# The multicellular incoherent feedforward loop motif generates spatial patterns

**DOI:** 10.1101/579342

**Authors:** Marcos Rodríguez Regueira, Jesús Daza García, Alfonso Rodríguez-Patón Aradas

## Abstract

The multicellular incoherent feedforward loop (mIFFL) is an extension of the traditional intracellular IFFL gene motif where the interacting nodes no longer need to be genes inside the same cell but can be spatially distributed in different cells. We studied for the first time the spatial computing abilities of these mIFFL through in silico simulations done with individual-based models (run in Morpheus and GRO software). We observed that: 1) a genetic circuit working as a mIFFL can behaves as an edge detector of the border of an infection, and 2) a mIFFL can be the inner mechanism generating the complex 7 stripe pattern of eve in D. melanogaster embryogenesis. So, in this work, we show that multicellular IFFL architectures can produce spatial patterns and are a promising spatial computing motif that deserves to be incorporated into the toolbox of pattern generation and multicellular coordination mechanisms. This study opens several future lines of research: multi-agent IFFL applied in ecology as a tool to predict spatial position of interacting animals or in distributed robotics.

## INTRODUCTION

### Patterning in Systems and Synthetic Biology

Patterning is a common area of study in Synthetic and Systems biology. At its most basic level, patterning can be defined as the process that leads to the features of an organism being arranged into a structured and ordered configuration. Such patterns can be spatial, temporal or both and may arise from many diverse underlying mechanisms [Kondo, 2011], such as the implementation of biological circuits based on recurrent architectures, better known as motifs [Alon, 2007; Schaerli, 2014]. These motifs exhibit well-known interactions that drive the dynamical behavior of the circuit. A combination of motifs with morphogen gradients has been used for spatiotemporal pattern formation [Basu, 2005; Tabor, 2009].

As the research trend has recently changed into multicellular environments [Amos, 2014; Kolar, 2015; Solé, 2016], in this work, we propose adaptions of network motives for their implementation in multicellular configurations. The use of spatially distributed motives [Macía, 2016] reduces the metabolic burden of the carrier and ease the implementation of the circuit, since each cell only runs a specific part of the circuit. Besides, the disposition of the different elements along the circuit, interconnected by diverse communication systems, will lead to a richer spectrum of patterns.

### Incoherent Feedforward Loops

The feedforward loops (FFL) are a type of motif where a factor, Z, is regulated under two different paths. On the one hand, there is a direct regulation from X to Z that defines the X-Z pathway. On the other hand, there is an indirect pathway (X-Y-Z) where a factor X regulates Z through a third element, Y. Whenever these two regulation paths are consistent, meaning that both activate or repress, then the FFL is coherent, when opposite, it is incoherent.

Depending on the combination of interactions, there is a classification of types of FFL architectures [Alon, 2007], represented in Fig. 1A. Additionally, feedforward motifs exhibit a dynamical behavior highly dependent on the architecture, on the strength of the interactions and on the time response of each component [Mangan, 2003]. They are widely present in nature in many biological regulatory networks. A good example of how a network, with an asymmetrical response as mentioned before, can become a reliable regulator is the Gal-system in *E. coli*, explained as a feedforward regulatory network in [Kaplan, 2008].

**Figure 1.**
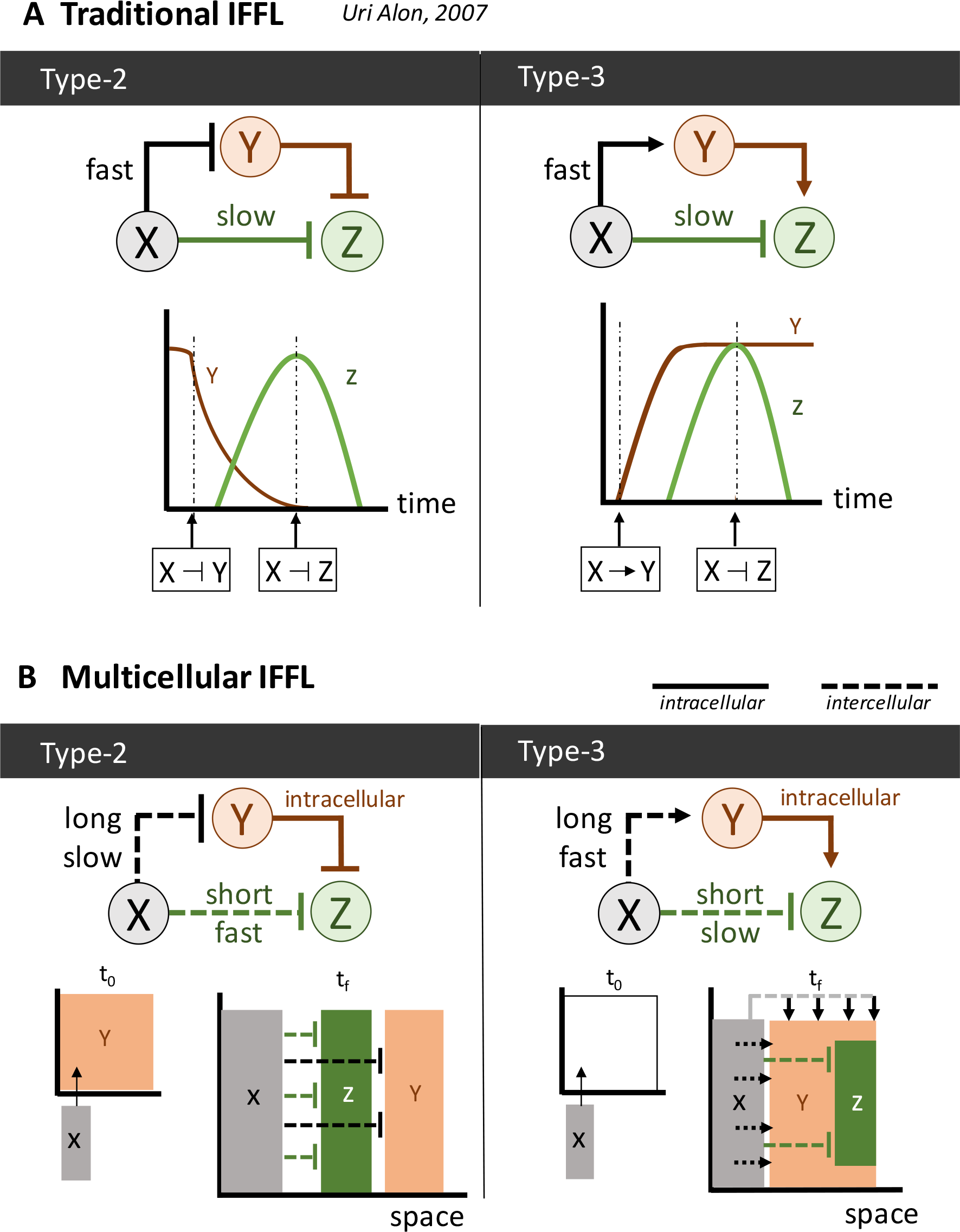
The Multicellular Incoherent Feedforward Loop, an extension of the traditional Incoherent Feedforward Loop. A) We present two possible IFFL configurations that can generate pulses of Z expression over time. In the type-2 IFFL, an input of X represses Y (expressed because it had no repressor) and this causes the activation of Z. When the slow repression from X to Z starts, then the expression level of decays. In the type-3 IFFL, as all the interactions are activations, the pulse of Z appears because of the difference in speed responses to an input of X. B) Explanation of the novel network motif proposed in this paper. The main change is that the interactions are now intercellular. At left, we show a multicellular type 2 – IFFL with particular conditions of speed response and spatial range. Whenever an input of X appears in some region of the space, it represses Y remotely in all the range of the black signal. Afterwards, the green signal inhibits the expression of Z in a shorter range, so a stripe of Z will be expressed between these two lengths. At right, we present the type3-mIFFL, where an input of X over blank space will activate Y in a region as large as the range of the black signal. Afterwards, the slower and shorter green signal inhibits the expression of Z and it is reduced to the terminal region of Y (its activator).

**Figure 1.**
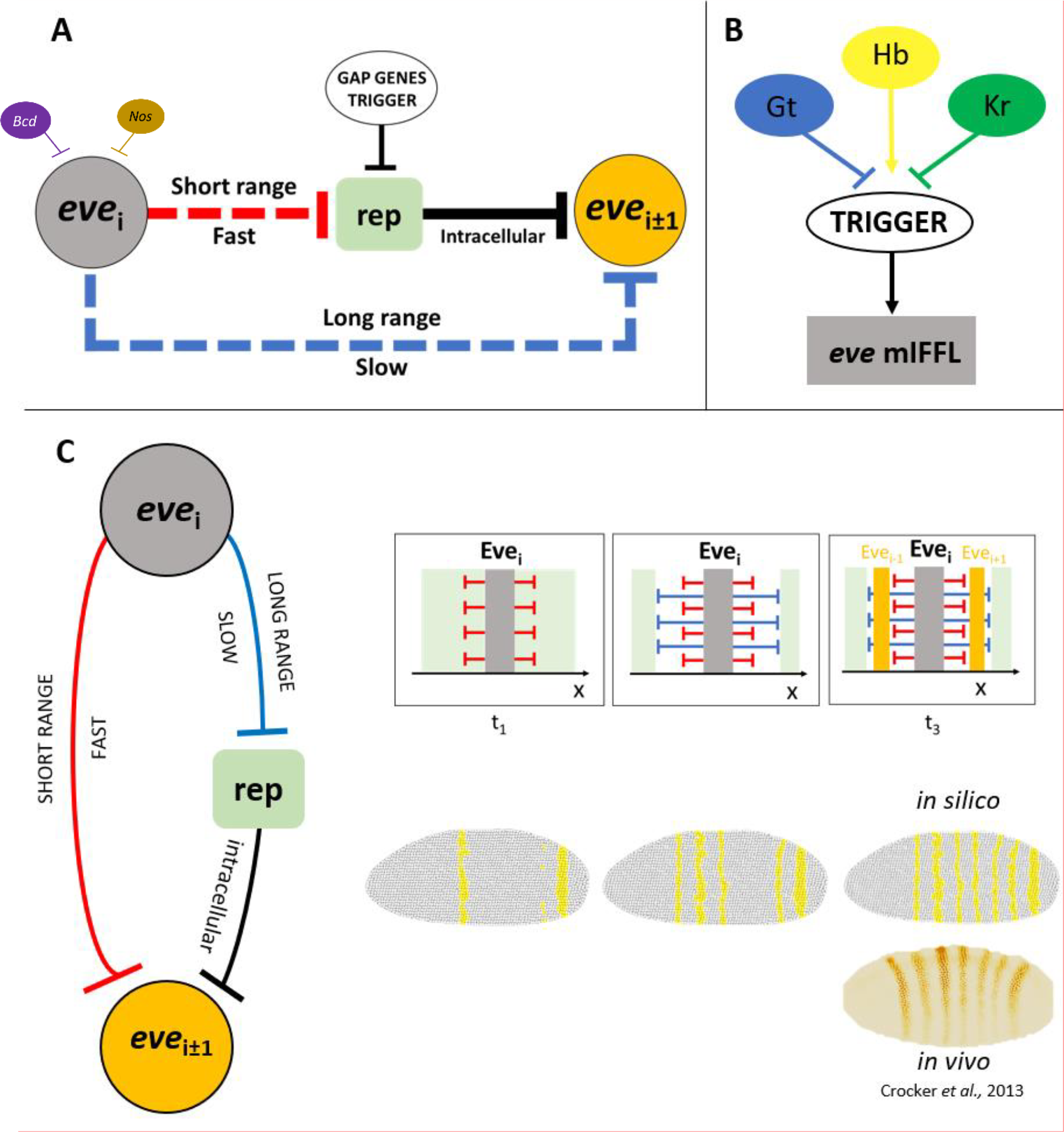
MIFFL motif for Eve pattern formation. A) Global mechanism of formation of the Eve stripes. The circuit is in Off state by default, because the repressor is activated along the embryo. This repressor is silenced in certain regions where the conditions of the gap gene trigger are met. These regions will be the initiation points of the pattern. Once Eve stripes are formed, they emit two signals that incoherently regulate the appearance of the next stripe. The first signal is fast and short ranged (red) and it prevents the activation of Eve near an existent band. The second signal (blue) is slower and long ranged and it represses the constitutive repressor, thus activates Eve expression. Bcd and Nos stop eve expression near the anterior and posterior region, respectively. The pattern becomes stablished when the Eve gene reaches the steady state both in space and time along the embryo. B) The gap gene trigger is a global enhancer of the mechanism. It switches the mIFFL state to ON in the regions where the Gt, Kr and Hb conditions are met. In the literature, these have been explained as the conditions for the formation of Eve2, but in our model, they also enhance Eve7. C) Formation of the Eve stripes during time. The model develops over time following a sequential formation of bands. The first ones appearing are EVE2 and EVE7. The neighboring bands appear after the incoherent regulation of the two signals over space following the logic detailed in Fig4A. Besides, we compare the final in silico pattern with an in vivo eve phenotype. It can be seen that the result is very similar.

**Figure 2.**
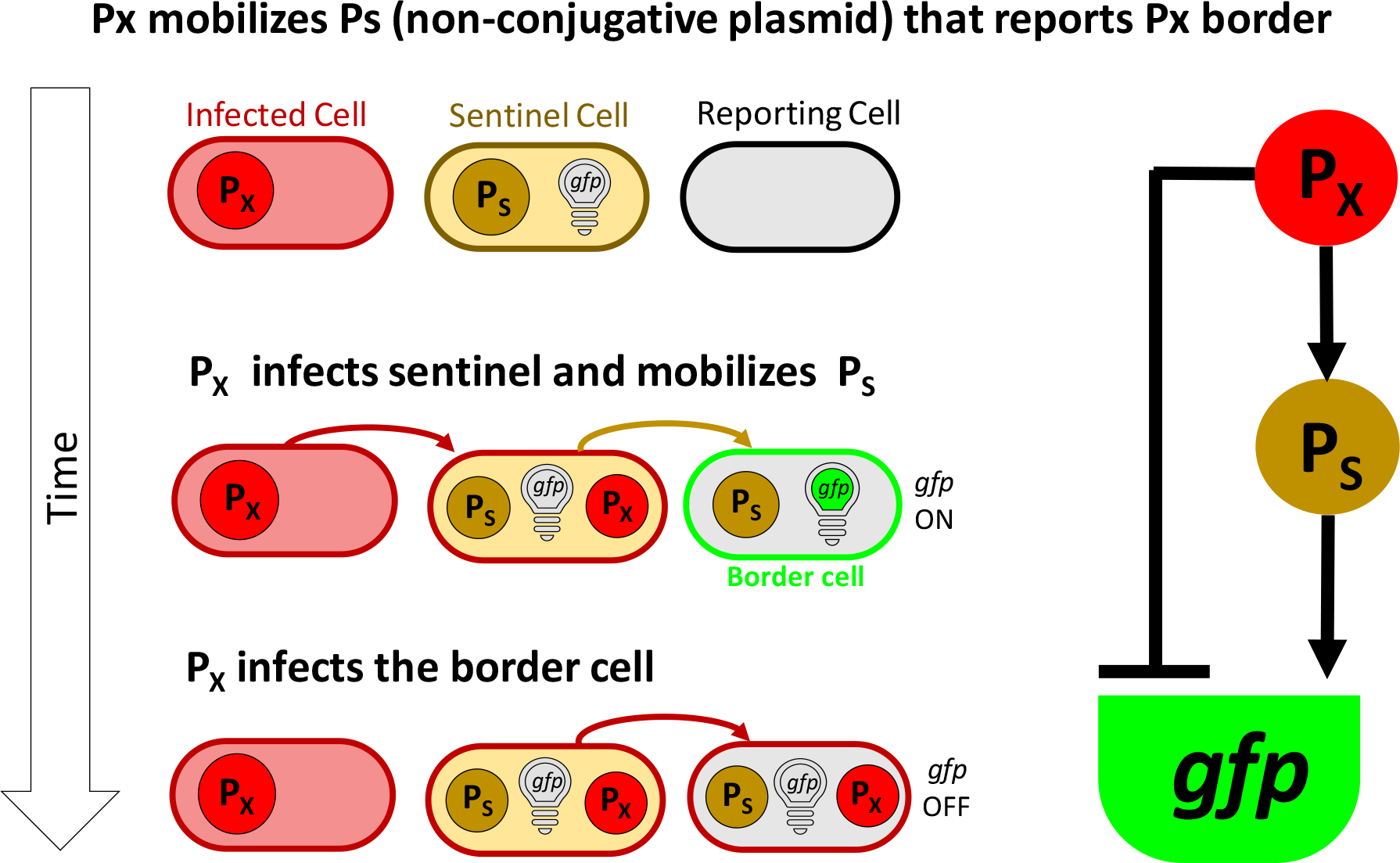
Edge detection logic. In this circuit there is an infective and self conjugative plasmid (Px), a sentinel and conditionally-conjugative plasmid (Ps) and a non-conjugative plasmid (Ps_0_), initially located in the same bacteria as Ps. The goal of the circuit is that the plasmid Ps arrives to a plasmid-free cell and then express the fluorescent protein GFP to indicate that this cell is in the outer border of the infection. The initial cells carrying Ps constitutively repress gfp expression by means of Ps_0_, avoiding that sentinel cells report the edge everywhere. The temporal evolution of the circuit is shown at left. 1) The infection plasmid conjugates to a sentinel cell. When both are together, the conditional conjugation of Ps gets activated and it can mobilize. Afterwards, it conjugates to a plasmid-free cell. This cell lacks the constitutive repression of Ps0, so gfp is expressed and the edge is reported. 2) When Px arrives to this border cell via conjugation, it represses gfp again. As it can be seen, Px incoherently regulates gfp, both directly (intracellularly) and indirectly, by means of the conditional conjugation of Ps. At, right we present the mIFFL architecture following this logic.

**Figure 2.**
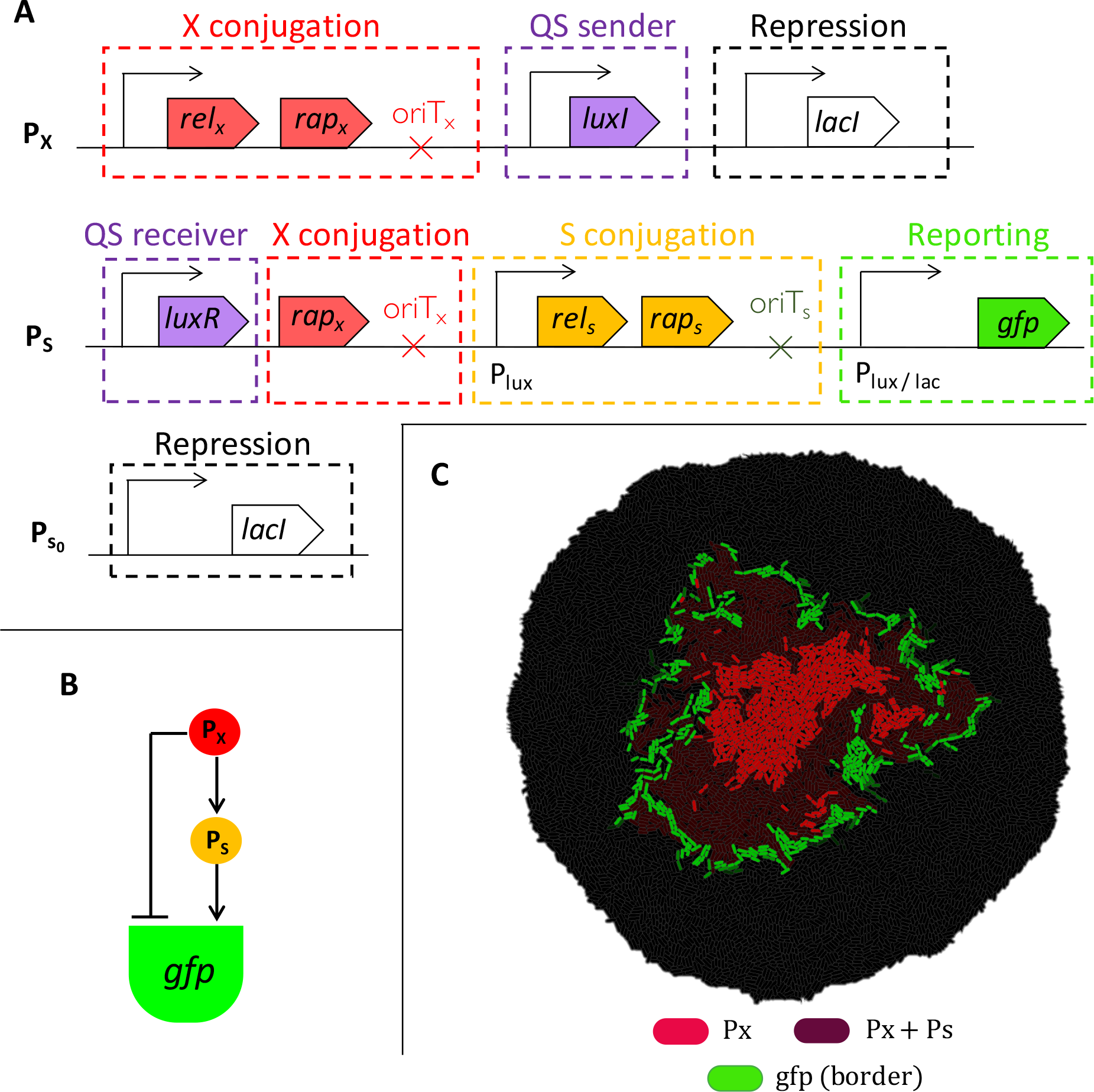
Edge detection system. A) Circuit design: There are three different plasmids present in the colony: Px, Ps and Ps0. Initially there are both plasmid-free cells (empty cells) and Px holders (infected cells). Afterwards, we introduce sensor cells, carrying both Ps (sentinel plasmid) and Ps0 (repressor plasmid). Our goal is to detect the border of spread area of Px, a self-conjugative plasmid, that constitutively expresses LacI and LuxI. Ps0 plasmid is not conjugative and can be found in initial sensor cells. Its function is the repression of gfp in Ps. Ps (sentinel plasmid) is conjugative under two different conjugative machineries associated to oriTx and oriTs. When Px is in its neighborhood, the diffusible autoinducer AHL (emitted by cells holding Px) and the protein LuxR bind and activate Plux, inducing the function of conjugative machinery associated to oriTs. In addition to that, Ps shares part of the conjugative machinery of Px, so when a coinfection occurs, Ps competes against Px for its relaxase, making Ps mobilizable through oriTx and, therefore, even more likely to be conjugated. The expression of gfp is regulated by a dual promoter Lux/Lac [Ayukawa, 2010] so it will only be expressed when the Ps plasmid is the only one in the cell (i.e. no lacI repressor is expressed in the cell) and the bacterium is in an area of high AHL concentration. B) IFFL motif of the design: The gfp present in the sensing plasmid is incoherently regulated by Px. GFP spatial spreading is induced by the self-conjugative plasmid Px by activating the conditional conjugation of Ps. When this plasmid is conjugated to an empty bacterium, then it reports gfp. Although, when the infection arrives to this same bacterium, then it is the plasmid Px, with lacI constitutively expressed, which represses the protein. c) Performance of the system: Typical pattern obtained in the simulations for an optimal set of parameter values and 300 min of simulation. Light red bacteria represent the infection; dark red represents coinfected bacteria (both sentinel and infection); light green bacteria are reporting the edge; black bacteria represent empty cells and repressed sentinels.

In this study, we will focus on the incoherent feedforward loops (IFFLs). It has been shown how these kind of incoherent networks form nonlinear behaviors like temporal pulses [Hart, 2013; Schaerli, 2014] and low/high pass filters [Sohka, 2009].

In some configurations of a traditional FFL, the components X, Y, Z are intracellular and interactions between them produce a temporal pulse of Z expression (Schaerli, 2014). The hypothesis of our work is that by using some intercellular interactions instead of their intracellular counterparts (bacterial conjugation or diffusible signals, for instance), then, the expected output will be a stripe-shaped spatial pattern. In the intracellular version of the circuit, different response times for each of the two regulatory paths are required for the stripe to appear. In the multicellular IFFL, this condition translates into intercellular interactions of different ranges. Further details about this novel design are addressed in Fig 1b.

In this work we have chosen an *in silico* approach, so the methods are the simulation software used, specifically, two individual based models (IBM). An IBM is a class of computational model that explicitly simulate the actions and interactions of autonomous agents. The IBMs have been used in many fields, like ecology. However, its use in Systems and Synthetic biology has begun recently. The IBMs are often composed of numerous agents, which represent each of the individuals. In our case, those agents are the cells of a synthetic bacterial colony and of a Drosophila embryo. The IBM is also composed of a decision-making heuristic, which characterizes the response of each individual of the system. For our model these decision rules will be, for instance, the probability of conjugation to a neighbor, in the case of *E. coli*, or the level of production of a certain morphogen by the cell, in response to certain concentrations of other morphogen, in the case of *D. melanogaster*.

We used the gro IBM [Jang, 2012; Gutierrez, 2017] for the simulation and evaluation of the *E. coli* synthetic circuit presented in this work. In gro, each cell contains proteins and plasmids. The plasmids determine which proteins are produced and those proteins, depending on its presence or absence, determine the behavior of the cell, including intercellular interactions. The behavior of the population then emerges from these interactions. Additionally, each agent can execute actions under certain conditions. An example of these actions is the light emission of the agent by a fluorescent protein expression, cell death, bacterial conjugation or environmental signal emission and absorption. This system can simulate a large amount of cells (in the order of 10^5^ bacteria) in less than 10 minutes. As gro also implements several forms of intercellular communication, it is ideal for prototyping and testing multicellular IFFL circuits on a large scale. We will focus mainly on two intercellular communication actions: conjugation and diffusible molecular signals.

The triggering of conjugation in gro is based on a frequency rate, where each cell carrying a conjugative plasmid is checked for the appropriate protein condition. If this condition is met, then, based on the conjugation rate, a probability is calculated for transferring the plasmid. Also, a recipient neighbor is chosen randomly. Conjugation is used as a programmable local communication mechanism and is a key participant in the designs presented in this paper.

On the other hand, the simulation of diffusible signals is based on bacterial quorum sensing. Sender bacteria emit a molecule to the environment and receiver bacteria are able to detect and absorb it. This way, one bacterium can change the genetic state of a far neighbor.

### Morpheus

Morpheus is the IBM selected for the *in silico* approach of *Drosophila melanogaster* embryogenesis. Morpheus is a modeling and simulation environment for the study of multiscale and multicellular systems developed at the University of Dresden [Starruss, 2014].

Morpheus allows defining models with different groups of cells and designing genetic networks that link them. Our model includes relations between different morphogenes and the activities of them within each cell. This is a suitable IBM because it allows for both intercellular and intracellular modeling. Another characteristic of Morpheus is that it is capable to solve partial differential equations, so it is able to estimate the concentrations of morphogenes at each point of the embryo. This capability is essential because the activations/inhibitions of the genes are driven by these concentrations.

Finally, both simulators have a graphical interface that allows the visualization of the results in a simple and comprehensible way by representing each cell individually.

## RESULTS

In this section we explain the mechanisms driving the proposed genetic circuits in this work and we show the results from the simulations. In the first place, we present a bacterial edge detection system, with the simulator gro. In the second place, we reproduce the stripped pattern of eve gene in *D.melanogaster* with the Morpheus simulator.

### A synthetic edge detection system

We present an edge detection circuit able to remark the border of an infective self-conjugative plasmid (Px) spreading area. The edge (i.e. the spatial pattern) arises from the interaction of this input plasmid with a conditionally mobilizable sensor plasmid, henceforth called infection and sentinel plasmids, respectively. By adding bacteria carrying a sentinel plasmid (Ps) to a colony where the infective plasmid proliferating radially outwards from a spot source, then the expected behavior of the system is to dynamically report the outer border of the area where bacteria with the infective plasmid are located.

In this document, the modularization of IFFL gene regulatory networks into different plasmids distributed along the colony is proposed for the dynamical generation of patterns. It is in this spirit that conditional conjugation is presented as a control mechanism over the horizontal spread of plasmids. Conditional conjugation is the process by which the transcription of the conjugative machinery of a plasmid is regulated. In this case, the plasmid will only be conjugative when the inducer/repressor of its expression is present/absent. In Fig.S1, we show a simple circuit where the conjugation of a plasmid is regulated by the presence of an autoinducer (AHL). In the presented edge detection system, the conjugation of Px into a cell carrying Ps induces its mobilization. Thus, conditional conjugation is crucial for our goal to be achieved, because we do not want the conjugation of Ps to occur randomly, but to be selectively guided within the colony.

Along with the conditional conjugation outcome, plasmids may also initiate intercellular communication actions with other parts of the distributed circuit through the emission of diffusible molecules. Small molecules and horizontal gene transfer are thus the two mechanisms used in the presented circuits for communication between different plasmids and forming the distributed motifs. The difference between both mechanisms makes the combination stronger, because plasmids spread by local contact between bacteria. Nevertheless, diffusible signals exhibit a higher diffusive rate and the capability of accumulation in case of excess of emitters [Ortiz, 2012; Goñi-Moreno, 2013].

In our edge detector, there is an initial colony with cells both empty and carrying the infective plasmid and. The sentinel plasmids (able to sense and report the infection) are inserted in new bacteria added to the colony. The reporting consists of the expression of the luminescence protein GFP. Initially, there is an intrinsic repression of this protein in the cell by means of an extra non-conjugative plasmid ps0. Otherwise, sentinels would report the edge everywhere. Here is where the previously explained concept of conditional conjugation becomes crucial.

When Px is spreading and arrives to a sentinel cell, then the conjugative machinery of Ps switches to an active state. After, the conjugation of Ps to a plasmid-free cell enables *gfp* to be expressed since there is no intrinsic repression of the gene. Whenever Px arrives to this same cell, it represses the *gfp* expression again, since this bacterium is no longer located in the outer border but instead belongs to the infected region. This happens due to the underlying incoherent regulation of *gfp* by Px. The indirect side of the motif induces its expression by mobilizing Ps, but then when Px reaches this point by conjugation, it represses the promoter regulating *gfp*. A visual explanation of the logic can be consulted in Fig2.

In summary, there are two spatially-separated pathways for the regulation of *gfp*: either Ps senses the proximity of the infection and it can conjugate and express *gfp*, or the infection finally arrives to a reporter cell and represses it. Therefore, the system can be considered a distributed multicellular type-3 IFFL that dynamically generates a spatial pattern consisting of the perimeter of the spreading area of Px (see figure Fig. 3B).

In our simulations, we found some issues in the design. These issues are independent of the plasmids logic, but momentous for the performance. One of them is the requirement for Ps to conjugate faster than Px. This is necessary for the spatial stripe to be generated because he indirect pathway of the mIFFL must be the fastest. As we want to determine the outer border of a region, it is necessary that the activator of the reporter arrives before the inhibitor.

The strategies to overcome it are based on the current knowledge about the origin of transfer (oriT) of plasmids [Del Campo, 2012]. By using a RP4-based plasmids as sensors and a R388 class as infection plasmids, we will get a noticeable difference in conjugation rates, since RP4 has a higher frequency of conjugation. This can be improved by including both the oriT of RP4 and the R388 one into the sequence of the sentinels. In case of coinfection, there will be a competition for the same conjugative machinery and it will reduce the velocity of spreading of Px.

Another critical issue is the use of quorum sensing as an intercellular channel. The version of this circuit that relies only on conjugation for communication showed poor results. Hence, an extra intercellular mechanism was included for Ps to notice when Px is nearby. In this case, the mobilization of the plasmid is regulated by a dual promoter, inducible by both an intracellular element of Px and a quorum sensing signal emitted by far infected cells. The infective plasmid is supposed to emit an autoinducer that allows Ps to sense its closeness in advance. This new feature leads the system to a much better performance.

Both features together with the initial logic of plasmids make the edge detection system very efficient. The SBOL diagram of the plasmids used in the high-performance edge detector are introduced in Fig. 3A

The final issue we identified is related to the distribution of sensing and empty bacteria used to detect the edge. Since the reporting module needs to be hosted by initially empty bacteria, if the amount of sentinels surrounding the infection is large, then the conjugation of the plasmid will be blocked due to the entry exclusion mechanism [Smillie, 2010], leaving no possibility for reporting. Further details of the importance of these phenomena are shown in SI2 and a sensitivity analysis about the ratios between the different types of plasmids and the effect on the pattern obtained can be found in SI2.

The edge detector simulation shown in Fig2c results from the inclusion of the totality of features enumerated above and the use of optimal parameters. As it can be seen, the final outcome is definitely successful in relation to the expected pattern.

Another interesting variant of the edge detection system has also been tested. It is based on an infection plasmid that can be arrested by the sensor in case of coinfection. In this case, the multicellular IFFL incorporates an extra inhibition path, so it can be considered as an even more extended version of the concept of mIFFL (more info at Fig. SI3).

In order to prove the versatility, scalability and robustness of the mIFFL as a pattern generator in bacterial colonies we tested two extra circuits, which are considerably different in behavior to the edge detector. At first, we present a spatial XOR system, different from previous designs that only focused on computation [Tamsir, 2011; Ji, 2013; Goñi, 2013]. This is an extended design of the edge detector (using two self-conjugative plasmids as input signals) able to detect the different regions of the space where these inputs are present: the 1-1 (both inputs present), the 1-0 or 0-1 (only one of them is present) and 0-0 (both are absent). The initial setup consists of two spot sources of the plasmids, that grow radially outwards. A more complex sentinel plasmid is added and the expected behavior is to report the edges of the regions mentioned above. Further details of the system and the results of a simulation are shown in Fig. SI4a.

Finally, we designed a band detection system able to report the area between two thresholds of an external inducer (IPTG). The pattern that arises is a centered ring that reports the area between two thresholds of IPTG concentration. If the concentration profile of IPTG remains constant, the shape of the ring will be constant. Nevertheless, the system can become dynamic if the IPTG concentration changes in time, thus changing the shape of the ring. As a big difference from the previous design of [Basu, 2005], in this case there two different type of plasmids with a part of the circuit. Cells are able to communicate with each other, runs the same circuit, thus reducing their metabolic burden. A detailed explanation of the plasmids and the results from a simulation are shown in Fig. SI4b.

In conclusion, the synthetic genetic circuits introduced in this paper show how the mIFFL is a versatile mechanism for the generation of functional multicellular biocircuits. We introduced an edge detection system, we extended it to two different inputs and finally we designed a spatially distributed version of a band detector. These circuits can also be seen as pattern generators in bacterial colonies as they form ordered structures over space. By adding new features to improve the results of the first version we obtained better performance in function and higher quality in the spatial arrangement.

### 7-stripe Eve pattern formation during Drosophila melanogaster embryogenesis

The embryogenesis of the fruit fly (*Drosophila melanogaster*) consists of the group of processes that control the transformation of a single cell into a mature individual of *Drosophila melanogaster*. During the early stages of embryogenesis two axes are defined, the dorso-ventral and the anterior-posterior [Kimelman, 2011]. For the definition of these axes, it is essential the action of the morphogenes, mobile molecules whose non-uniform distribution drives the formation of differentiated structures.

These genes are sequentially expressed and drive the embryo to the differentiation of three regions (head, thorax and abdomen) [Gilbert, 2001]. The effects of the morphogenes in the formation of the dorsoventral axis has been previously studied [Hart, 2013]. Besides, they induce the segmentation of these zones.

One of the genes involved in this process is *even-skipped* (*eve*). Eve belongs to a gene group called pair-rule genes and plays a crucial role in the formation of the 14 segments of *D. melanogaster*. However, for the expression of this gene, it is necessary the presence of other groups of genes such as the maternal-effect genes and the gap genes. Both groups are expressed before the pair-rule genes.

The maternal-effect genes have a non-homogeneous distribution prior to the formation of the embryo [Macdonald, 1996]. This non-homogeneous distribution has its origins in the oocyte and it is caused by the differential affinity of the maternal-effect genes over the microtubules of the oocyte [Cha, 2001]. The most relevant representative of this group is *bicoid* (*bcd*). These morphogenes do not only have a direct effect over the pair-rule genes (including *eve*), usually acting as transcriptional activators, but also regulate the expression of the gap genes [Driever, 1990].

The expression of the gap genes marks the beginning of the embryo segmentation process, which starts during the cell cycle number 13. The interactions between these genes are quite complex and some are not clear yet. Nevertheless, the most relevant interactions are well understood [Jaeger, 2011]. The outcome of this network is the formation of a characteristic pattern of expression for every morphogen [Struhl, 1992] that will be the basis for the next group of genes to begin their expression, the pair-rule genes.

There also exists another group of genes that regulates the expression of the gap and pair-rule genes, the terminal genes. This group is only expressed in the anterior and posterior ends of the embryo. The terminal genes play a major role in the formation of structures such as the head or the tail and they normally repress other genes in these zones [Janssens, 2013]. The most relevant gene of this group is *tailless* (*tll*) that is expressed both in the anterior and posterior ends.

The pair-rule genes begin to express once the patterns of the gap genes are stablished towards the end of the cell cycle 13. There is a total of 8 pair-rule gene [Schroeder, 2011] and they give rise to very characteristic patterns of expression, consisting on thick stripes arranged perpendicularly to the anterior-posterior axis.

*Even-skipped* is one of the most studied pair-rule gene as it is the first of this group that is expressed and have influence in the rest pair-rule genes [Clark, 2017]. *Eve* originates a pattern of expression composed of 7 bands, perpendicularly arranged to the anterior-posterior axis. The current explanation by a series of independent enhancers, each of them activating a cluster of bands [Hare, 2008; Harding, 2018]. The mechanism of some enhancers has been widely studied, such as the one giving rise to the second stripe of Eve (Eve2) [Small, 1992; Goltsev, 2004; Bothma, 2014]. Other enhancers are not so well understood and even there are some stripes whose enhancers are still unknown. Even if the enhancer is known, it is extremely difficult to determine which morphogen activates or represses it. The combination of all these factors made extremely difficult the development of a global model that give rise to this pattern.

In order to stablish a global mechanism that explains the formation of all the *eve* bands, we develop an *in silico* model of an embryo of *D. melanogaster* in the beginning of the anterior-posterior segmentation, when the embryo is in the blastoderm stage between cell cycles 13 and 14. All the data related to the length, size, shape and number of cells of the embryo refer to this stage of development and can be found in SI5a. Moreover, we simulate the expression of the maternal-effect genes as an input and the gap-gene spatial pattern is dynamically generated from them, according to the interaction network shown in Fig.4A. Finally, the *eve* bands are obtained by means of an autoregulatory mIFFL. The spatial distribution of its expression will be very susceptible to both the concentrations of the maternal-effect genes used as input and the *gap-genes* obtained.

It is necessary to consider that at this stage of development of the embryo some genes have been already expressed so, at the beginning of the simulation, some morphogenes from the maternal-effect and gap-gene groups will be initially located. For these inputs, we used interpolation polynomials based on experimental data from [Perkins, 2006], that can be consulted at SI5b.

In order to define the dynamical concentration of morphogenes we use an SDD (Synthesis, Diffusion, Degradation) model [Ellis, 2009]. The diffusion and degradation parameters are defined as constants and their values, extracted from the literature. To define the synthesis of every morphogen, we make use of the Unc-logic model from [Perkins, 2006], based on logical rules.

In the first place, we simulate the formation of the pattern of the gap genes. With this simulation, we have obtained 2D images, and then we compare them to experimental images of *D. melanogaster* embryo. Simulations of the gap-genes expression are very similar to the *in vivo* results, as can be seen in Fig.4b.

Once the gap genes are fully expressed, the expression of pair-rule genes and, therefore, *eve* begins. *Eve* is expressed in 7 bands perpendicularly arranged to the anterior-posterior axis of the embryo.

In order to obtain this pattern in our model, we propose an mIFFL motif. In this motif Eve activates and represses itself by two independent and incoherent pathways. First, we introduce constitutive repressor that inhibits the expression of Eve along the embryo. We also propose that cells expressing Eve can emit two intercellular signals: a fast and short-range self-inhibitory signal and another slow, long-range signal that inactivates the constitutive repressor and, consequently, induces the expression of Eve. By means of this incoherent regulation and together with an adequate choice of parameters, we ensure that the new *eve* stripes appear in the desired position. The newly formed stripes have the same behavior, so they keep stablishing the pattern. Finally, to explain the absence of Eve stripes in the anterior part of the embryo, we take Bcd as a concentration-dependent repressor to prevent the expression of Eve in the anterior border.

Nevertheless, this system needs a trigger (see Fig 5A-B) to start because otherwise the constitutive repressor of Eve will never be silenced, and *eve* genes will never be expressed. So, there must exist an external mechanism that inhibits the constitutive repressor and switches the mIFFL on. As it has been previously discussed there are numerous enhancers of Eve stripes, but some of them are not yet well understood so we take as a trigger the best documented one, the one that activates the expression of Eve 2 and Eve 7. We have chosen this enhancer because it is clear which morphogenes activate and inhibit it, but there could be more starting points apart from this. We also obtain promising results with the simulations of the mutants for some of the morphogenes, this fact sets a positive evidence of taking the enhancer of the stripes 2 and 7 as a trigger to this mechanism (SI7).

**Figure 4.**
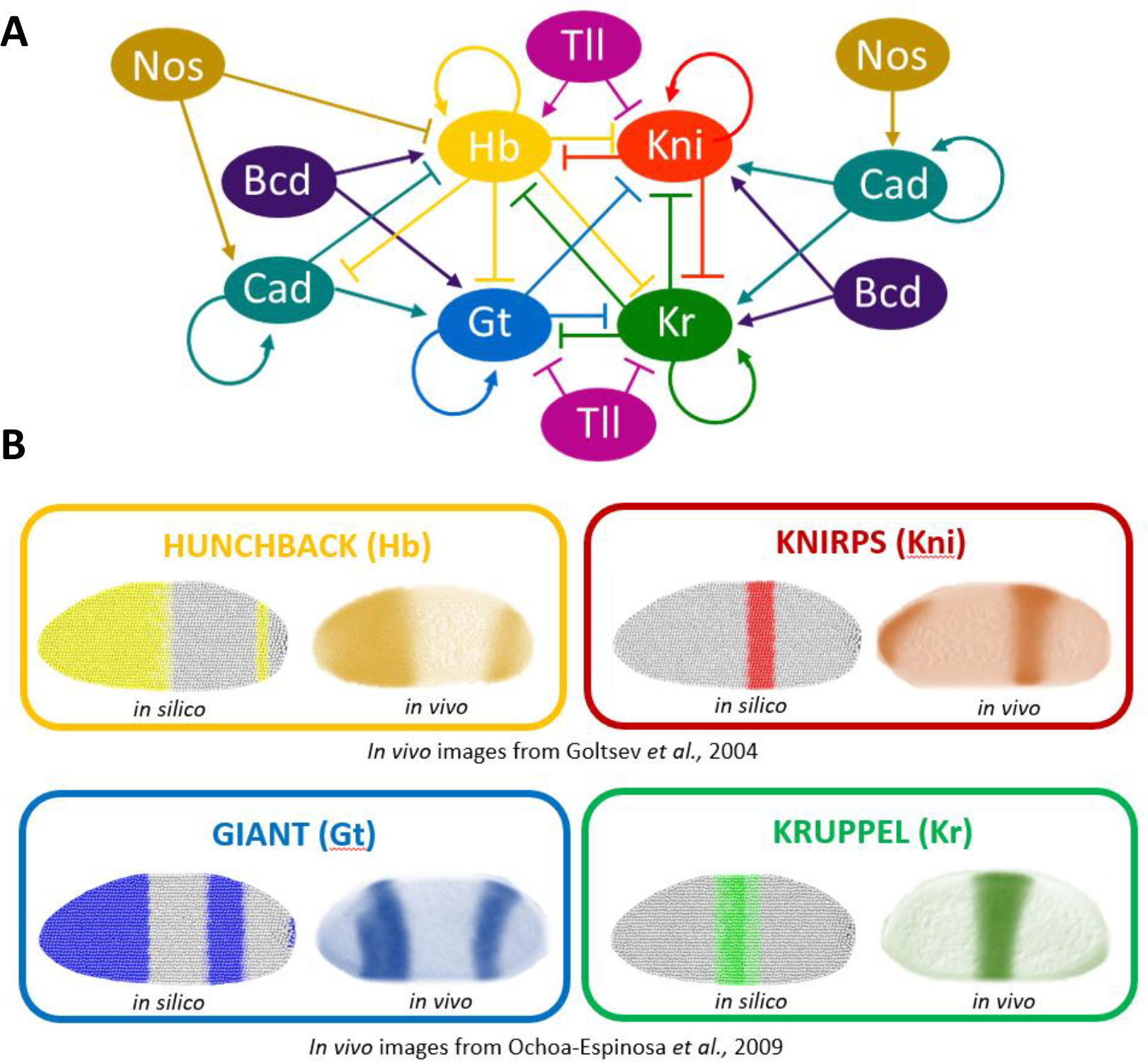
Gap Gene Network. A) Interaction network of the gap genes and maternal-effect genes. It is still unclear the strength of all these interactions. B) Comparison between in vivo and in silico gap-gene expression regions. Hb presents a full expression from the anterior end to the center and a smaller band near the posterior border. In the simulations, we reproduced these behaviors. Kr is expressed in a central band, although this band is wider in the in vivo results. Also, in both cases, Gt is expressed in two bands bordering the central region. In our model we have expression of giant in the anterior end region. This could be due our model does not simulate most of the terminal genes, a group of genes which is only expressed in the terminal regions and with a great influence there. Finally, Kni is correctly expressed in our model in the central region but it lacks the small anterior band. This also could be caused by the lack of many terminal genes in our model.

The simulations of Eve expression are very similar to the *in vivo* results. We obtain the same number of stripes and also these stripes have a similar wide and space between them compared to the experimental observations (see Fig 5C).

The result of this simulation supports our hypothesis that with an mIFFL it is possible to generate the *eve* pattern in *D. melanogaster*.

## CONCLUSIONS

Traditional 3-gene IFFLs motifs were developed to model intracellular genetic networks. In this work we have extended the IFFL motif to multicellular environments. In a multicellular IFFL the interacting nodes are no longer genes inside the same cell. The nodes can be spatially distributed over space and not necessarily in the same cell. The interaction between nodes is done by intercellular communication.

With the *in silico* simulations done with IBMs (Morpheus and GRO) we have shown that: 1) a mIFFL can work as an edge detector of the border of an infection, and 2) a mIFFL can generate the 7 stripe pattern of even-skipped in D. melanogaster embryogenesis. So, in this study, we have probed that type-2 and type-3 multicellular IFFL architectures are able to produce spatial patterns. Why are these precise IFFL motifs able to process spatial information? The main reason is that the indirect arms of the motif are supposed to be longer ranged than the direct ones, so there is a spatial delay in the mobilization of elements that drives to dynamical spatial patterns.

This study, that introduces the mIFFL motif for spatial computing, opens several promising future lines of research:

- It is an open problem for future research whether other types of multicellular incoherent (or coherent) feedforward loops are also robust strategies for spatial information processing.
- mIFFLs could be applied to model other processes of cellular differentiation in embryogenesis and developmental biology, as for example, the somitogenesis in vertebrates [Hester, 2011].
- The study of other variants of mIFFLs and mCFFLs that can also be interesting, not only for spatial patterning, but also as a tool to model higher-order interaction motifs, present in complex microbiomas, like human microbiota or for programming complex multicell consortia.
- The possible extension of mIFFLs to multi-agent IFFL in ecology, where the nodes would be the different species interacting in an ecosystem, would allow to predict the spatial distributions of the animals (analogously as done with “rock-paper-scissor” ecology models).
- In distributed robotics, multi-agent IFFL motifs could be used to program interactions between robots to generate complex spatial robot distributions.

## ACKNOWLEDGMENT

This work was supported by the European Union project PLASWIRES (612146/FP7-ICT-FET-Proactive), by Spanish MINECO projects TIN2016-81079-R and by the grant S2017/BMD-3691 InGEMICS-CM, funded by Comunidad de Madrid (Spain) and European Structural and Investment Funds.

## S1. CONDITIONAL CONJUGATION

**Figure S1.**
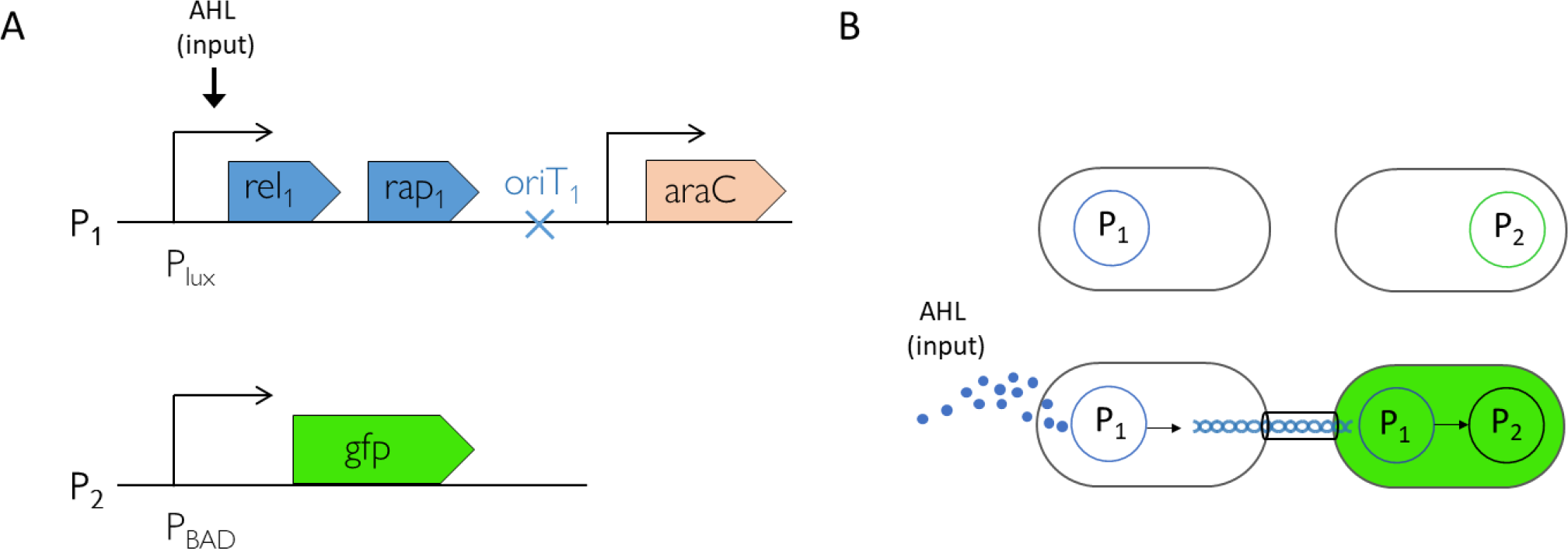
Scheme of the conditional conjugation process. The conjugative machinery of bacteria is formed by a *relaxase* (Rel) and the *relaxosome-accessory proteins* (Rap). These two proteins are essential for the conjugation process and are specific to the origin of transfer (oriT) of each plasmid, making it a highly orthogonal and scalable control mechanism [Smillie, 2010]. In this figure, we show a synthetic genetic circuit driven by the conditional conjugation process. A) P1 is a conditionally conjugative plasmid that constitutively expresses *AraC*, which is the activator of *gfp* in plasmid P2. This way, if we can control the mobility of P1, then we will be able to control the fluorescence of bacteria carrying P2. B) In this case, it is a high concentration of an autoinducer (AHL) what triggers the expression of the conjugative machinery (Rel and Rap) in P1. Whenever this plasmid arrives to a bacterium carrying P2, then *gfp* will be expressed and the fluorescence will be noticeable.

## S2. PARAMETER SWEEP ON EDGE DETECTOR

**Figure S2.**
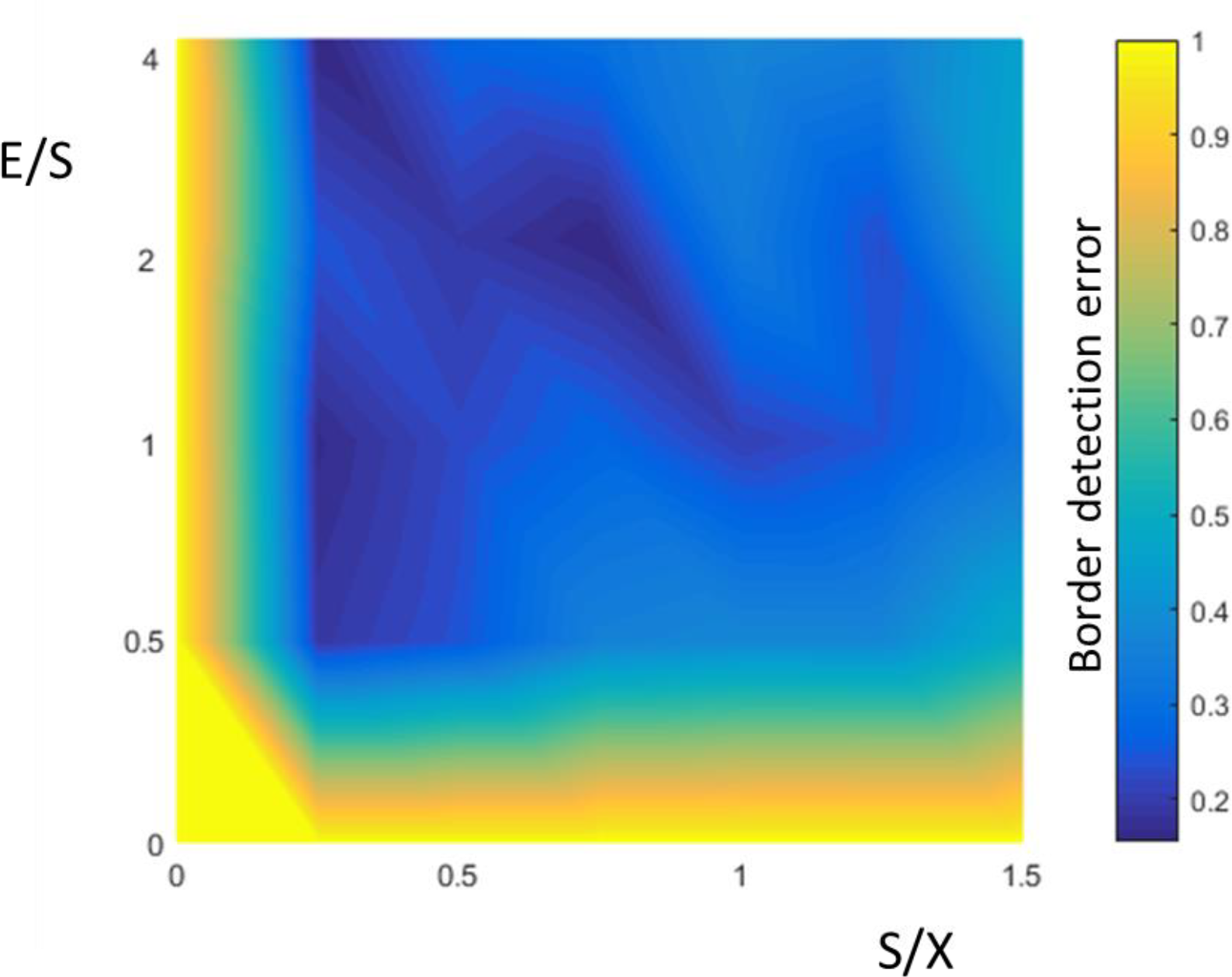
Parameter sweep on bacteria ratios. After all the improvements in the design, the system is still very susceptible to two parameters. These are the ratios between empty (E) and sensor (S) bacteria and between sensor (S) and infected (I) bacteria. Larger E/S parameters induce to less coinfection between Px and Ps. Whereas in the lower ratios almost all of the infected cells have been coinfected, as it increases, more small Px areas appear in the colony. Under these conditions, the reporting is worse, because an infection is needed to report. However, using small E/S ratios drives the colony to the blocking issue. It consists of sensor bacteria keeping the infected cells confined, so a lot of coinfections are needed for Ps to reach an empty cell. This makes the detection system slower, so it is necessary to establish an adequate ratio, which solves the blocking problem in a proper manner. In the case of S/I, high ratios drive to the blocking problem while low coinfection appears with low ratios. In order to search the optimal parameters, we make a heatmap representing both ratios in the axes and the error in edge detection (estimated with image processing morphological techniques [Marquez-Neila, 2014]). The parameters which best fit the requirements are E0/S0=2 and S0/I0=0.75.

## S3. VARIANT OF EDGE DETECTOR

**Figure S3.**
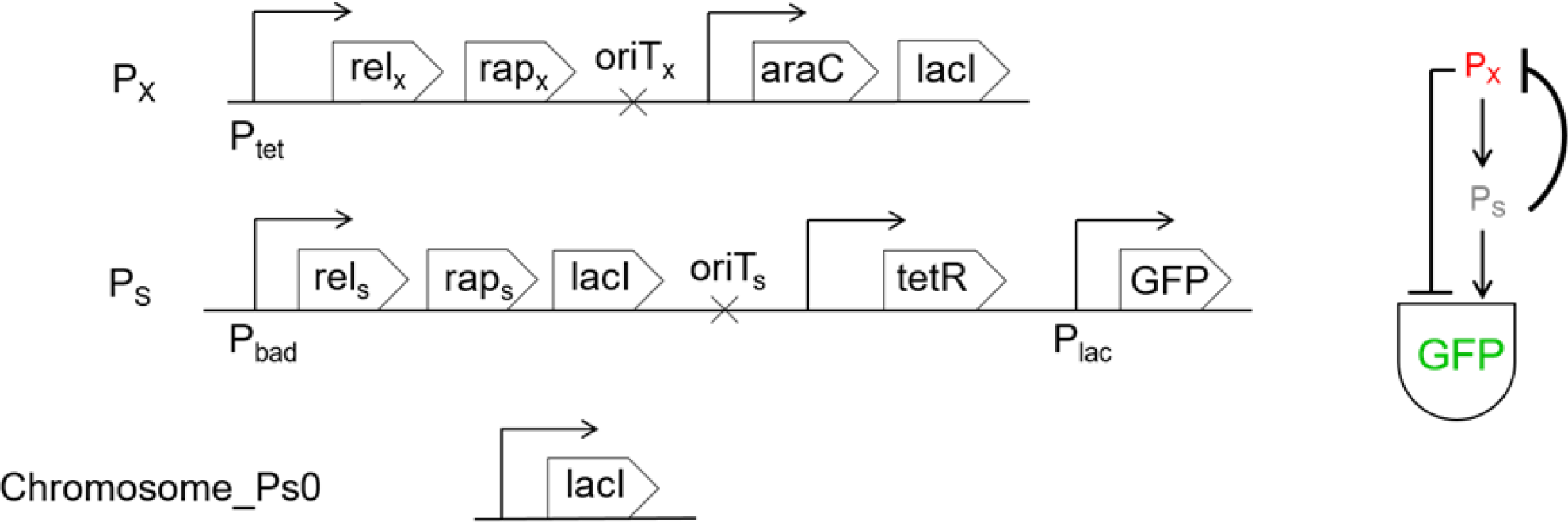
Edge detector with plasmid arresting. In this figure we present an alternative design of the edge detection system, where the sentinel plasmid not only reports the edge but also stops the conjugation of Px. In this case, both Px and Ps are conditionally conjugative, but Px is repressible and Ps is inducible. This feature makes the system more functional, because it prevents the spreading of the infection, but also more synthetic, because the infective plasmid has to be conditionally conjugative and regulated by a factor of Ps.

Furthermore, it shows an even more complex extension of mIFFLs, with feedback between nodes. This complex architectures yield very rich spatial behaviors.

## S4. XOR AND BAND DETECTOR

**Figure S4.1.**
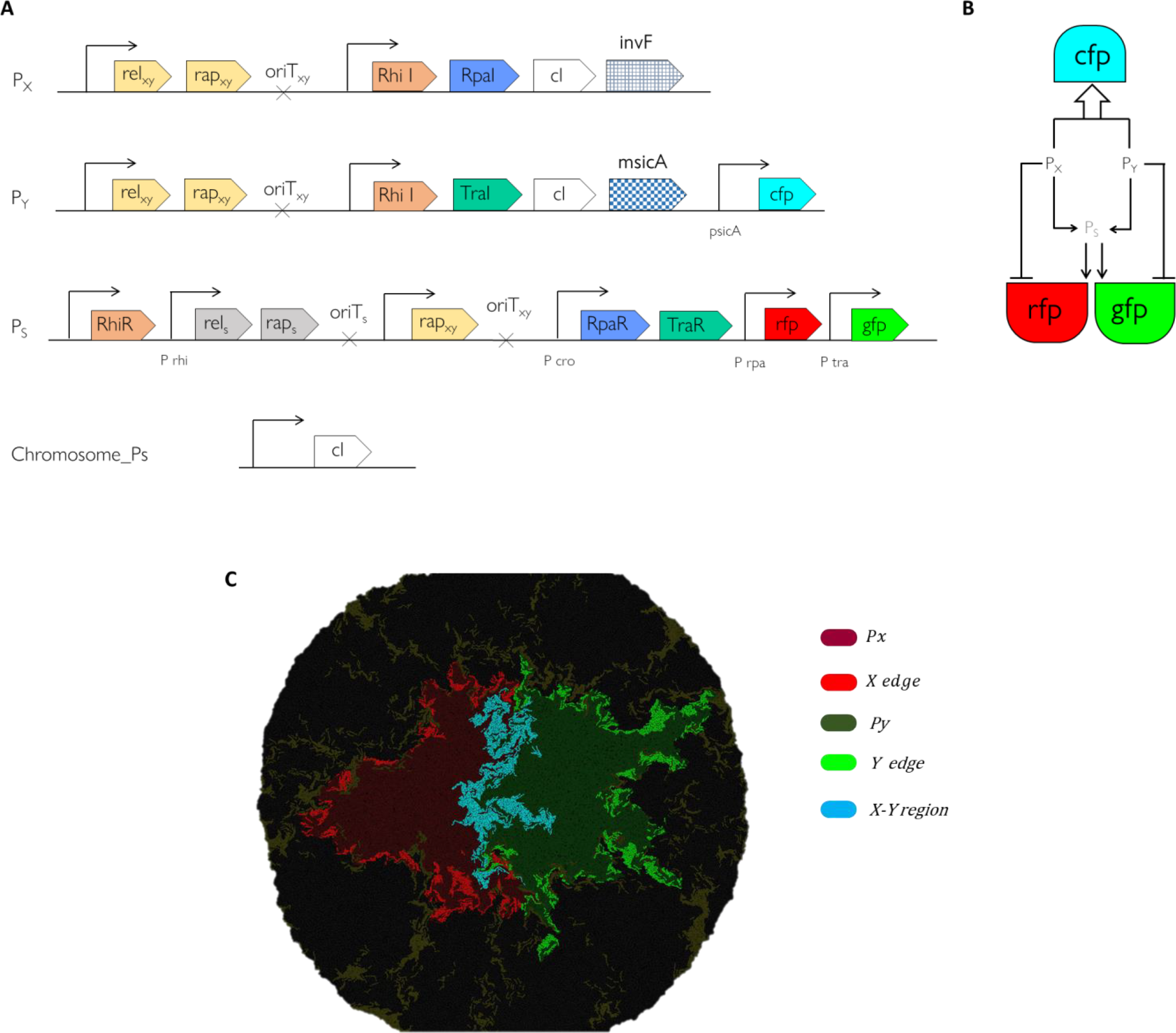
Spatial XOR from two coupled IFFLs. A) Circuit design: There are four different plasmids present in the colony: two self-conjugative input plasmids, Px and Py, which are initially placed in the center of the colony close to each other; a mobilizable sensor plasmid Ps and its accompanying Ps0, which are initially randomly distributed in the colony. Both input plasmids share their conjugative machinery. In the same way as the edge detector, both input plasmids emit a quorum sensing signal (Rhi), which is absorbed by bacteria holding Ps. Following the logic presented in the edge detector design, the conjugative machinery of Ps is activated when it senses that Px or Py are nearby. Equally, in this case, Ps lacks a conjugative protein from the conjugative machinery of the inputs. This way, when a coinfection occurs, Ps competes against Px or Py for this protein, enhancing the conjugation probabilities of the sensing plasmid even more. On the other hand, the reporting module is regulated by other quorum sensing signals that are completely orthogonal, according to [Scott, 2016]. Cells carrying Px and Py, emit Tra and Rpa, respectively, inducing the expression of the associated fluorescent proteins (green and red respectively) in sensor bacteria. In the edge detector design, the expression of the reporter protein only occurs when the sensor protein is in an empty cell where no *cI* is expressed. In order to differentiate the 1-1 region, *cfp* is expressed under the presence of both input plasmids. Each input plasmid contains either sequence for a part of a transcription factor (SicA* or its chaperone InvF). Both components couple when in presence of each other. This regulation mechanism is typical from Salmonella typhimurium and acts as an AND gate, so when both parts are present, the promoter PsicA is activated [Moon, 2014]. Finally, Ps0 plasmid is a non-conjugative plasmid that can be found in the initial cells bearing Ps. Its function is the repression of the reporting proteins in Ps. B) Design motif: Two independent IFFLs like the ones presented in the edge detector are coupled through a common node (Ps). In addition, there is a non-incoherent regulation of *cfp* reporting the presence of both inputs. C) Typical pattern obtained after 300 min of simulation and for the optimal set of parameters. Dark colored regions represent each source of infection while light colors represent each edge; the blue central area is the XOR 1-1 region and it also acts as an inner edge for each plasmid; dark yellow bacteria are the initial sensors.

**Figure S4.2.**
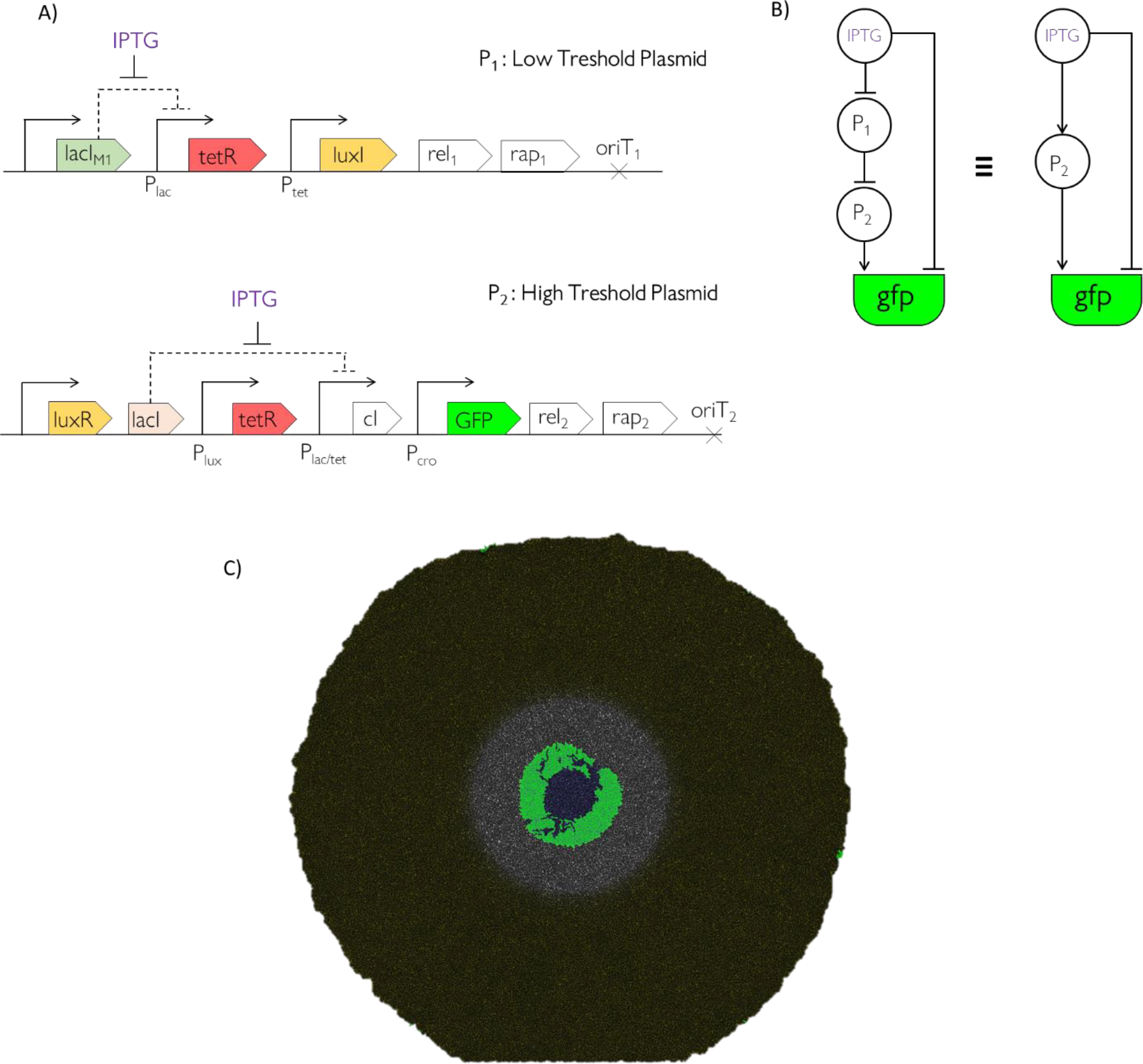
Band detector. A) Circuit design: There are two different plasmids present in the colony: P1 (low threshold) and P2 (high threshold). Both plasmids are regulated by a Plac promoter, so the concentration of IPTG represses the LacI protein in certain concentrations. The key of this design is the use of two Plac promoters sensitive to different concentration thresholds [Lakshmi, 2008]. Bacteria holding P2 start reporting gfp when the concentration is lower than the higher threshold. When the concentration is lower than the lower threshold, P1 will start preventing the reporting of P2. In sum, in the area where IPTG concentration is higher than the highest threshold, LacI will be expressed in both plasmids; in the area where LacI is silenced only in P2, gfp will be reported and, finally, in the area where both plasmids are activated a quorum sensing signal is emitted by cells with P1 and it represses gfp in P2. B) IFFL motif of the design: The gfp reporting gene is regulated by a type-3 IFFL architecture, where an IPTG source controls the expression and mobilization of the two plasmids P1 y P2. In the simplified version of the motif (presented in the right), IPTG activates P2 by means of AHL. The full motif includes the repression cascade from the input analog signal to P1 and the repression from P1 to P2. As there are two different IPTG concentration thresholds, the regulation will be distributed in separate areas. C) Pattern obtained for 120 min of simulation with a single central source of IPTG (gray and blue colors), so the two thresholds can be easily detected.

## S5. MATHEMATICAL MODEL OF DROSOPHILA EMBRYO

### 1 Geometry

The space where the simulations were carried out is a mask of a real embryo of *Drosophila Melanogaster*. The embryo is in the stage 6 of gastrulation (Interactive Fly), the number of cells at this stage is around 6000 (Zalokar, 1976). We only simulate 2200 because we are only representing the external surface of one lateral face of the embryo.

The size of the embryo at this stage is approximately 0.5mm long (Drocco, 2011)

**Figure S5.1.**
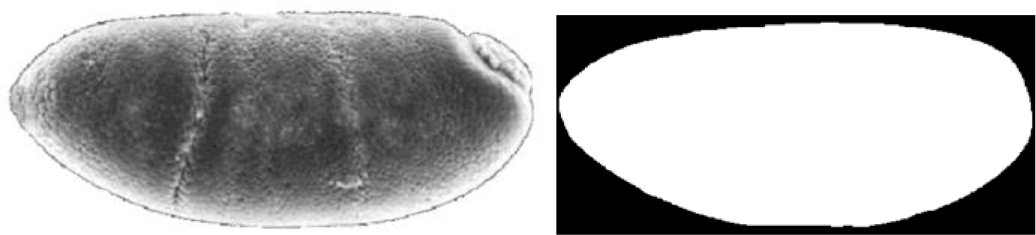
Embryo of *D. Melanogaster* and the mask derived. Gastrulating embryo - lateral view - stage 6. This mask was used for the simulations in Morpheus

### 2 PDE + Unc-Logic Model

The production and decay rates have units min^−1^, and the diffusion rates have units (1% embryo length)^2^ min^−1^

|  | Diffusion | Decay | Unc-Logic Rules | Production |
| --- | --- | --- | --- | --- |
| Hb | 2.16 | 0.146 | $((\text{Bcd} > 19 \text{ or } \text{Hb} > 49 \text{ or } \text{Tll} > 101) \text{ and } \text{Cad} < 122 \text{ and } \text{Kr} < 139 \text{ and } \text{ni} < \text{K4} \text{ and } \text{Nos} < 110)$ | 31.3 |
| Cad | 2.25 | 0.00243 | $((\text{Nos} > 125 \text{ or } \text{Cad} > 150) \text{ and } \text{Hb} < 61)$ | 7.65 |
| Kr | 0.67 | 0.067 | $((\text{Bcd} > 9 \text{ or } \text{Kr} > 103 \text{ or } \text{Cad} > 151) \text{ and } \text{Hb} < 177 \text{ and } \text{Gt} < 4 \text{ and } \text{Kni} < 115 \text{ and } \text{Tll} < 20)$ | 16.3 |
| Gt | 0.4 | 0.087 | $((\text{Bcd} > 35 \text{ or } \text{Cad} > 133 \text{ or } \text{Gt} > 85) \text{ and } \text{Hb} < 215 \text{ and } \text{Kr} < 15 \text{ and } \text{Tll} < 102)$ | 16.1 |
| Kni | 0.9 | 0.067 | $((\text{Bcd} > 5 \text{ or } \text{Cad} > 152 \text{ or } \text{Kni} > 94) \text{ and } \text{Hb} < 7 \text{ and } \text{Kr} < 141 \text{ and } \text{Gt} < 70 \text{ and } < 18)$ | 17.0 |
| Constitutive Repressor (Cons) | 0.0 | 1.0 | $(\text{LRS} < 0.1)$ | 1.0 |
| Short Range Signal (SRS) | 5.0 | 0.03 | $(\text{Hb} > 40 \text{ and } \text{Gt} < 30 \text{ and } \text{Kr} < 10 \text{ and } \text{time} > 40 \text{ or } (\text{Eve} > 14))$ | 50.0 |
| Long Range Signal (LRS) | 4.0 | 0.01 | $(\text{Hb} > 40 \text{ and } \text{Gt} < 30 \text{ and } \text{Kr} < 10 \text{ and } \text{time} > 40 \text{ or } (\text{Eve} > 14))$ | 50.0 |
| Eve | 0.0 | 0.0 | $(\text{Hb} > 40 \text{ and } \text{Gt} < 30 \text{ and } \text{Kr} < 10 \text{ and } \text{Nos} < 105 \text{ and } \text{Bcd} < 70 \text{ and } \text{time} > 40) \text{ or } (\text{SRS} < 0.1 \text{ and } \text{Cons} < 0.1 \text{ and } \text{Bcd} < 70 \text{ and } \text{Nos} < 105)$ | 2.0 |

The dynamic model of the genes is modeled by the synthesis, diffusion and degradation (SDD) equation (Grimm, 2010)

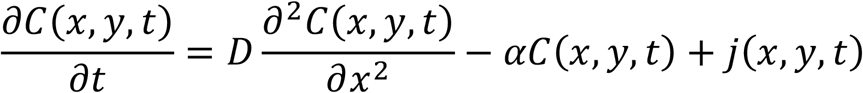

Where *C*(*x,y,t*) represents the concentration of a morphogen at time *t* and position *x,y*. *D* is the diffusion coefficient, α the degradation rate and *j*(*x,t*) describes the morphogen production.

### 3 Inputs

These are the polynomials of interpolation used as input for the *gap gene* simulation.

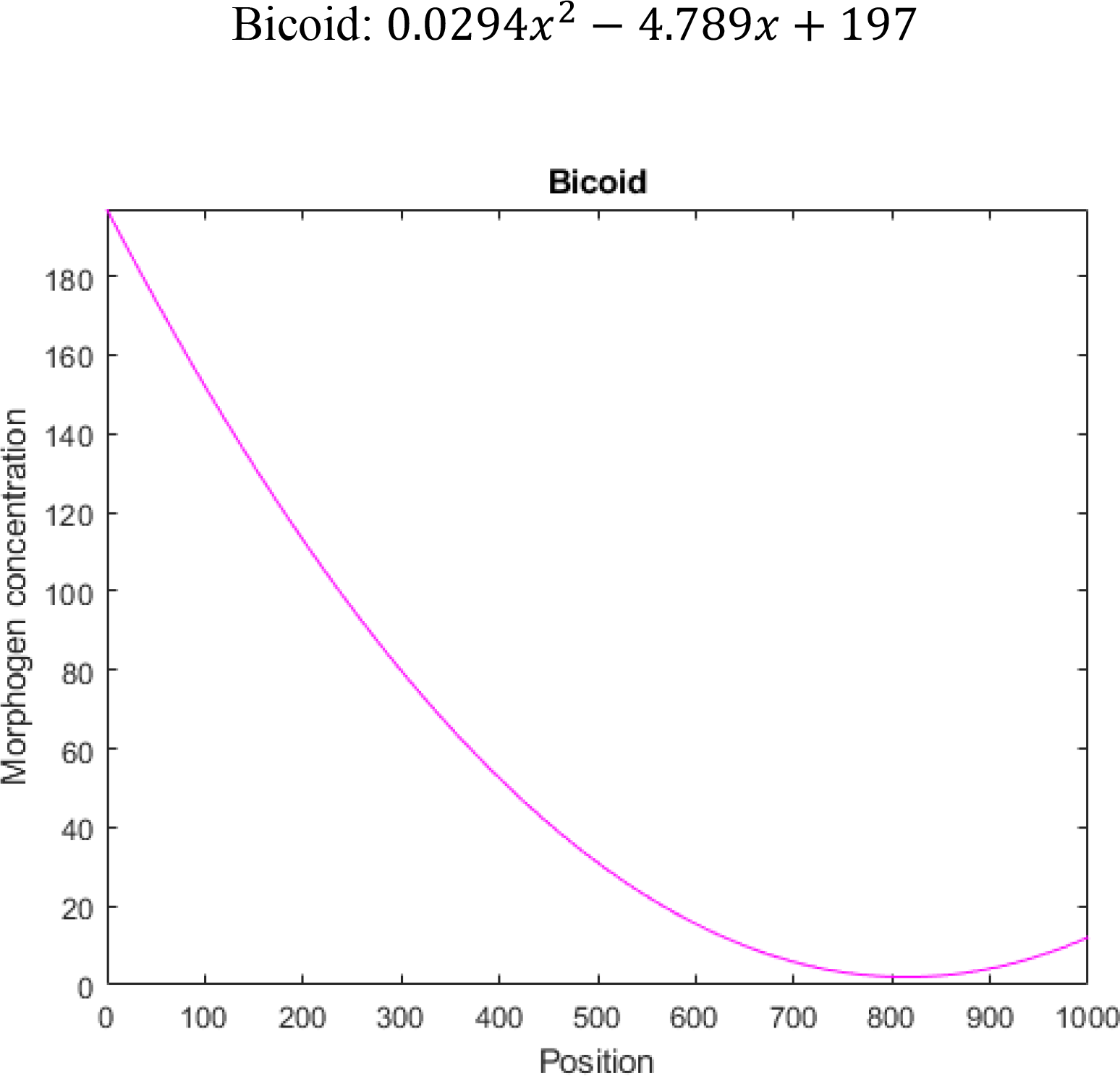

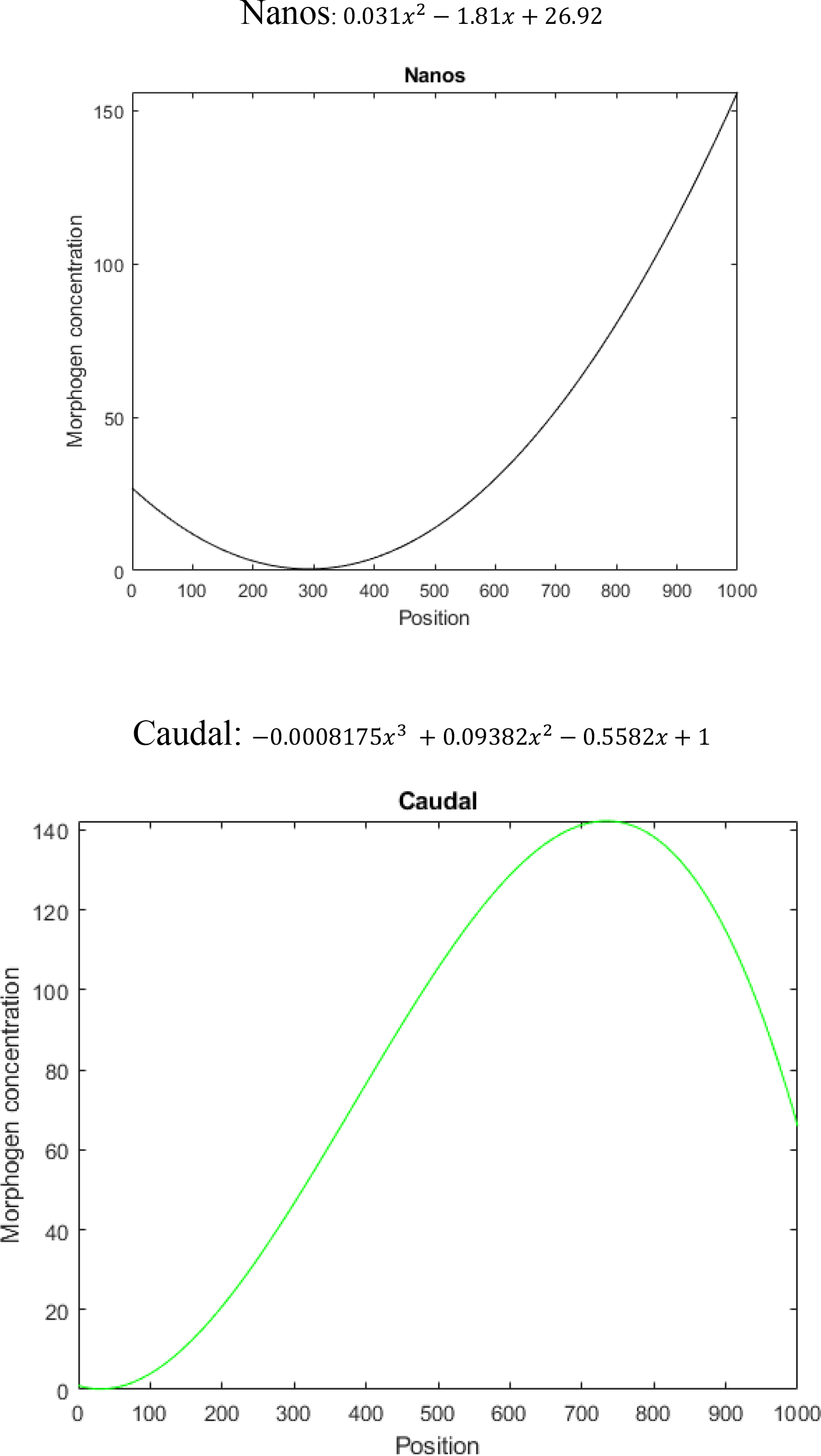

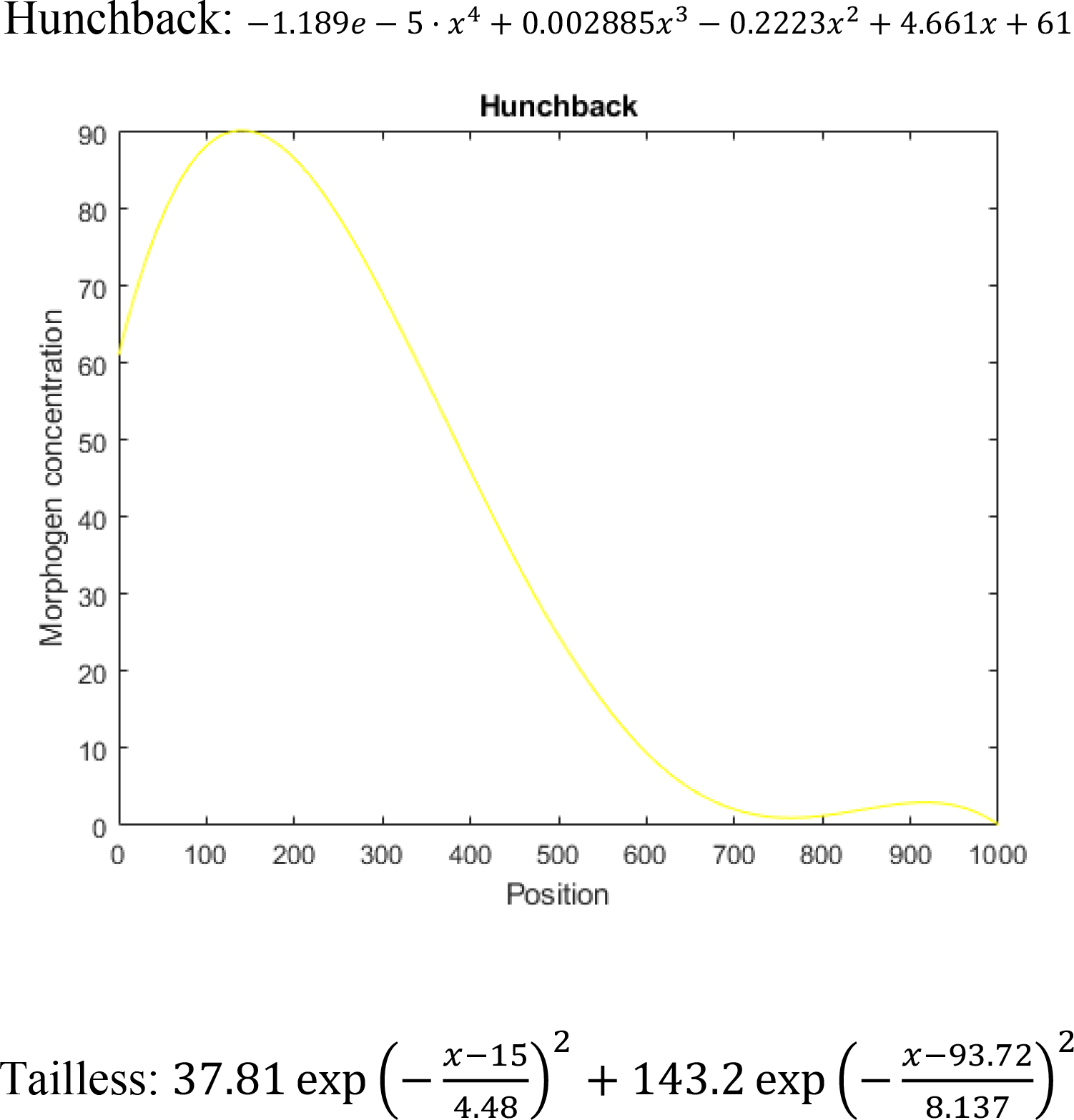

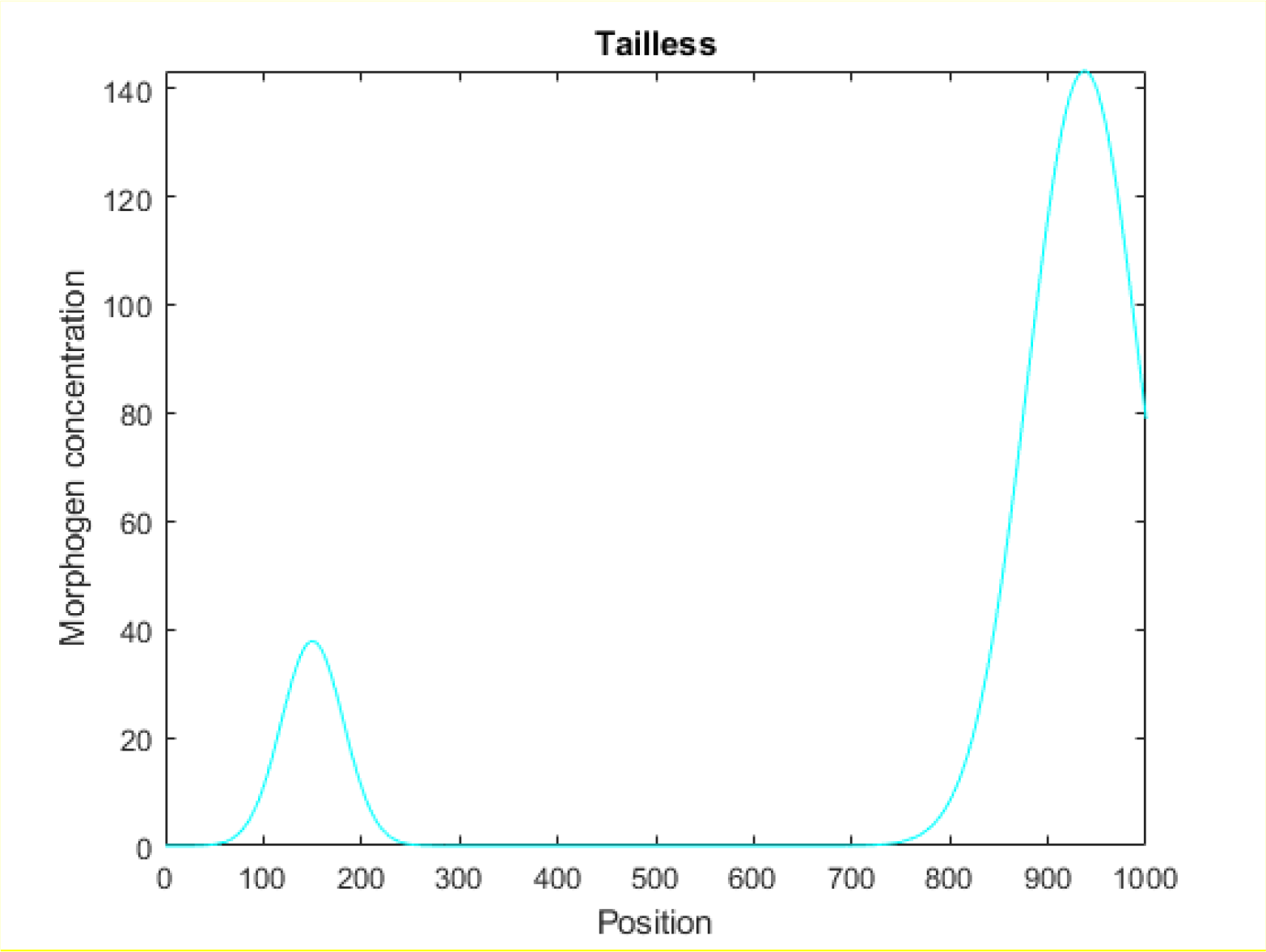

## S6. CONCENTRACIONS OF GAP GENES OVER SPACE

**Figure S6.**
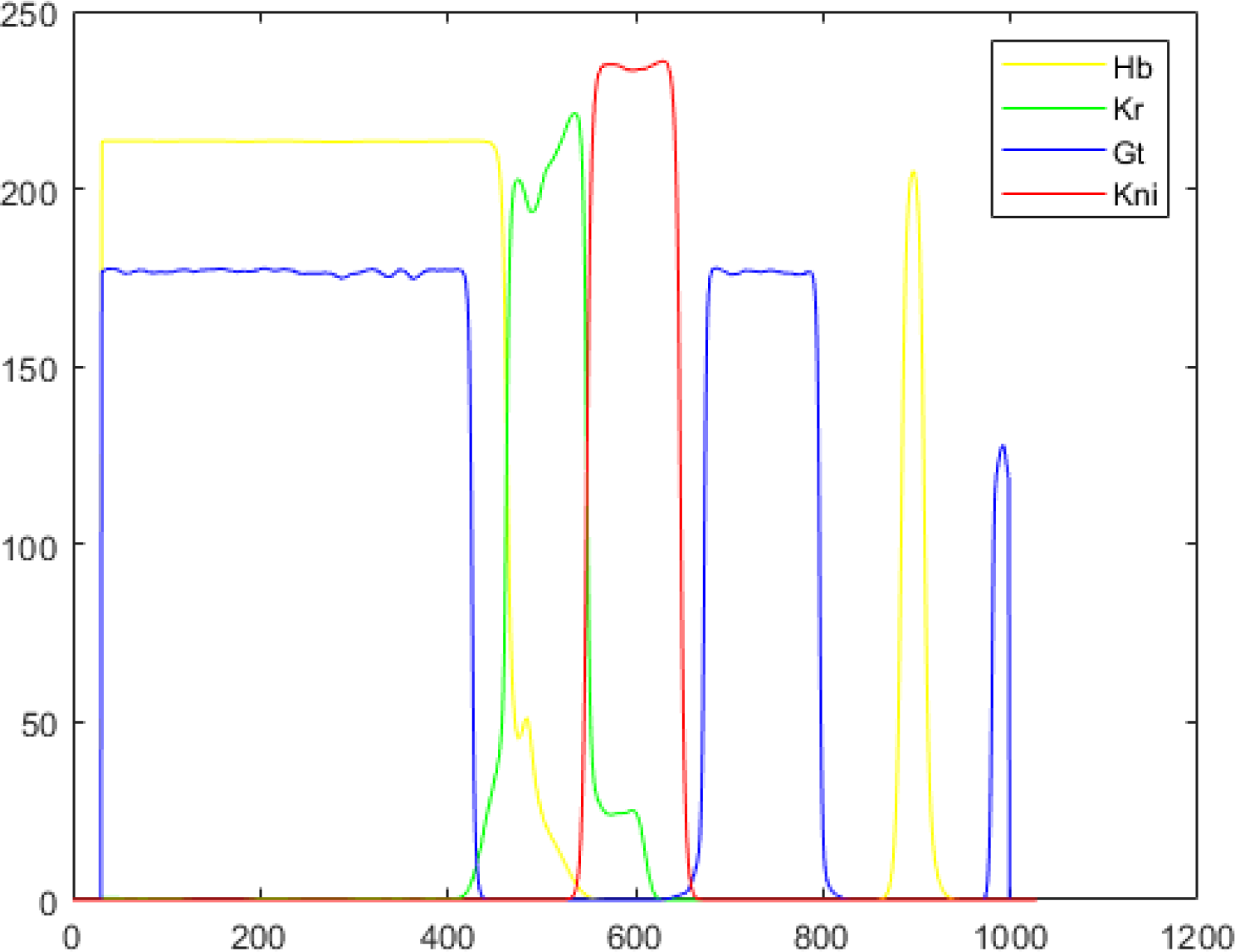
Concentration of Gap Genes. In this figure we show the spatial profiles of gap genes obtained after the simulation. These are the profiles after 40 min of simulation times and through an horizontal line in the center of the embryo. The results are very similar to the literature.

## S7. MUTANTS

In general, our model does not correctly reproduce the mutants patterns of expression of Eve. However, in some cases it partially reproduces some patterns of expression as it can be seen in the fig S7.

We will need to further analysis to refine the parameters.

**Figure S7.**
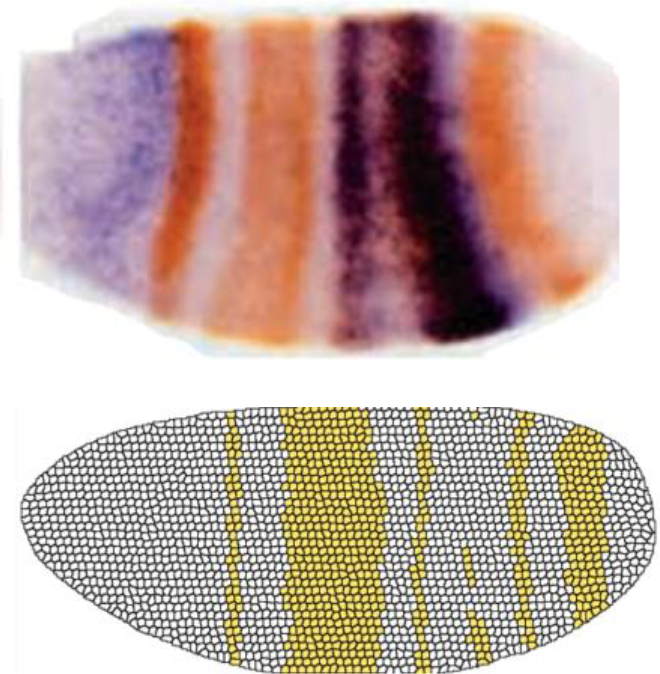
Comparison between in vivo and in silico gap-gene expression regions. The anterior part is correctly reproduced in our model with an overexpression of Eve2 that include Eve 3, nevertheless in the posterior region even though we correctly reproduced Eve 4 and Eve 7, Eve 6 and Eve 5 are not correctly reproduce.

